# Development of a multiplex immunofluorescence panel to study heterogenous cancer-associated fibroblast subtypes with spatial resolution

**DOI:** 10.64898/2026.06.26.734718

**Authors:** Amy Burley, Tatiany Silveira, Nick James, Manuel Salto-Tellez, Tom Lund, Anna Wilkins

**Affiliations:** Division of Radiotherapy and Imaging, Institute of Cancer Research, London, United Kingdom; Integrated Pathology Unit, Royal Marsden NHS Hospital Trust and the Institute of Cancer Research, London, United Kingdom; Royal Marsden NHS Hospital Trust, London, United Kingdom; The Precision Medicine Centre, Queen’s University Belfast, Belfast, BT9 7JU, United Kingdom

**Author notes:** **Corresponding author**: Anna Wilkins.

**Keywords:** Cancer-associated fibroblasts, single-plex immunofluorescence, multiplex immunofluorescence, spatial proteomics

## Abstract

**Background:** Single cell RNA sequencing provides a wealth of information to explore the complexities of the tumour microenvironment, but crucially the spatial topology of the tumour is lost and studying cellular interactions is limited. Spatial transcriptomics aims to address this however the technique remains cost prohibitive for the generation of data from meaningfully-sized clinical cohorts. In contrast, spatial proteomic profiling with multiplex immunofluorescence, preserves spatial interactions, is relatively cost accessible, and is scalable for large clinical cohorts to address powerful translational questions. Whilst multiplex approaches have advanced in recent years, we note that cancer-associated fibroblasts (CAFs) have been explored in less detail, potentially due to difficulties associated with CAF heterogeneity and the diversity of markers used to define them.

**Methods:** We designed, optimised, and validated a multiplex immunofluorescence panel that combines four frequently used CAF markers; alpha smooth muscle actin (αSMA), fibroblast activation protein (FAP), podoplanin (PDPN) and platelet-derived growth factor receptor alpha (PDGFRα) with CD8 and pan-cytokeratin. Here we share our methodology and the practical considerations taken to inform the final panel design. We also highlight the benefits of robust optimisation experiments.

## 1. Introduction

Cancer-associated fibroblasts (CAFs) are a heterogenous population of non-malignant cells that are abundant in the tumour microenvironment with functionally diverse roles that are often associated with promoting tumour growth, metastasis and immune evasion [1]. In contrast, some CAF subtypes have been associated with anti-tumour properties [2–4], as such targeting and reprogramming CAFs has become an attractive strategy to help overcome resistance to standard-of-care therapies.

Classification of CAF subtypes is a rapidly growing area of research and is essential to define the diversity within the population to ensure that CAF targeting strategies are effective and accurately identify CAFs with pro-tumour functions. Emerging studies suggest that CAFs from different tumour types have shared functional characteristics based on common biological principles [5]. Broadly, CAFs are defined as myofibroblast-like CAFs (myCAFs) that are associated with enhanced contractility, migration and extracellular matrix remodelling or inflammatory CAFs (iCAFs) that manipulate cells in the tumour microenvironment via secretion of inflammatory cytokines [6–9].

Cues from the local tumour microenvironment and communication with neighbouring cells contribute to the activation of CAFs and can influence the CAF subtype and functional status [10–12]. Moreover, interactions between CAFs and immune cells can contribute to immunosuppression and an immune excluded phenotype [13, 14], therefore preserving the spatial arrangement of the tumour microenvironment is critical to understanding how CAFs interact with other cells and the consequences this may have on treatment efficacy.

Historically, histological approaches to study CAFs have used a single marker such as alpha smooth muscle actin (αSMA) to act as a surrogate pan-CAF marker, however, one of the challenges of accurate identification and classification of CAFs is the lack of a specific CAF marker to distinguish them from other cell types and further characterise CAF subtypes. A multiplex approach enables the study of multiple markers in parallel to provide confidence in the classification of CAFs and gain insights into their functional state. By preserving the spatial integrity of tissue, analysis of multiplex images extends beyond quantification of cellular abundance and offers insights into the organisation of cellular neighbourhoods and spatially localised communication between CAFs and other cell types of interest.

Using many of the same principles as traditional immunohistochemistry (IHC), multiplex systems have been developed to facilitate the simultaneous study of multiple markers of interest, at protein or RNA level. Whilst the application of multiplex panels in a clinical setting may be restricted due to the cost and complexity of the analysis, for research purposes multiplexing can be a valuable tool by providing a wealth of information from a single section of tissue. This is particularly relevant when handling precious clinical trial material in limited supply.

High-plex technologies make use of metal and DNA-based antibody conjugation methods such as imaging mass cytometry [15] and the PhenoCycler (Quanterix) that facilitate the study of approximately 30-100 markers. Other newly-emerging platforms such as GeoMx, Visium, Visium HD and Xenium offer new approaches to spatially map transcriptomic data, ranging from a targeted panel of ∼500 genes to the whole transcriptome. Whilst technically impressive, it important to consider the balance between the number of markers and the number of samples that can be processed cost effectively within a reasonable timeframe [16].

We opted to use the 6-plex fluorescence-based Opal-tyramide signal amplification (TSA) system and PhenoImager scanner (formerly Vectra Polaris) from Quanterix (Figure 1A) for development of a multiplex immunofluorescence CAF panel. The system has a number of advantages: Firstly, opals can be paired with off-the-shelf antibodies and optimised into a working panel within a relatively quick time frame; Secondly, reagents can be used in existing autostainers to facilitate staining *en masse*, with a full 6-plex protocol complete overnight; Thirdly, scan times are relatively quick, resulting in a much higher sample throughput.

**Figure 1:**
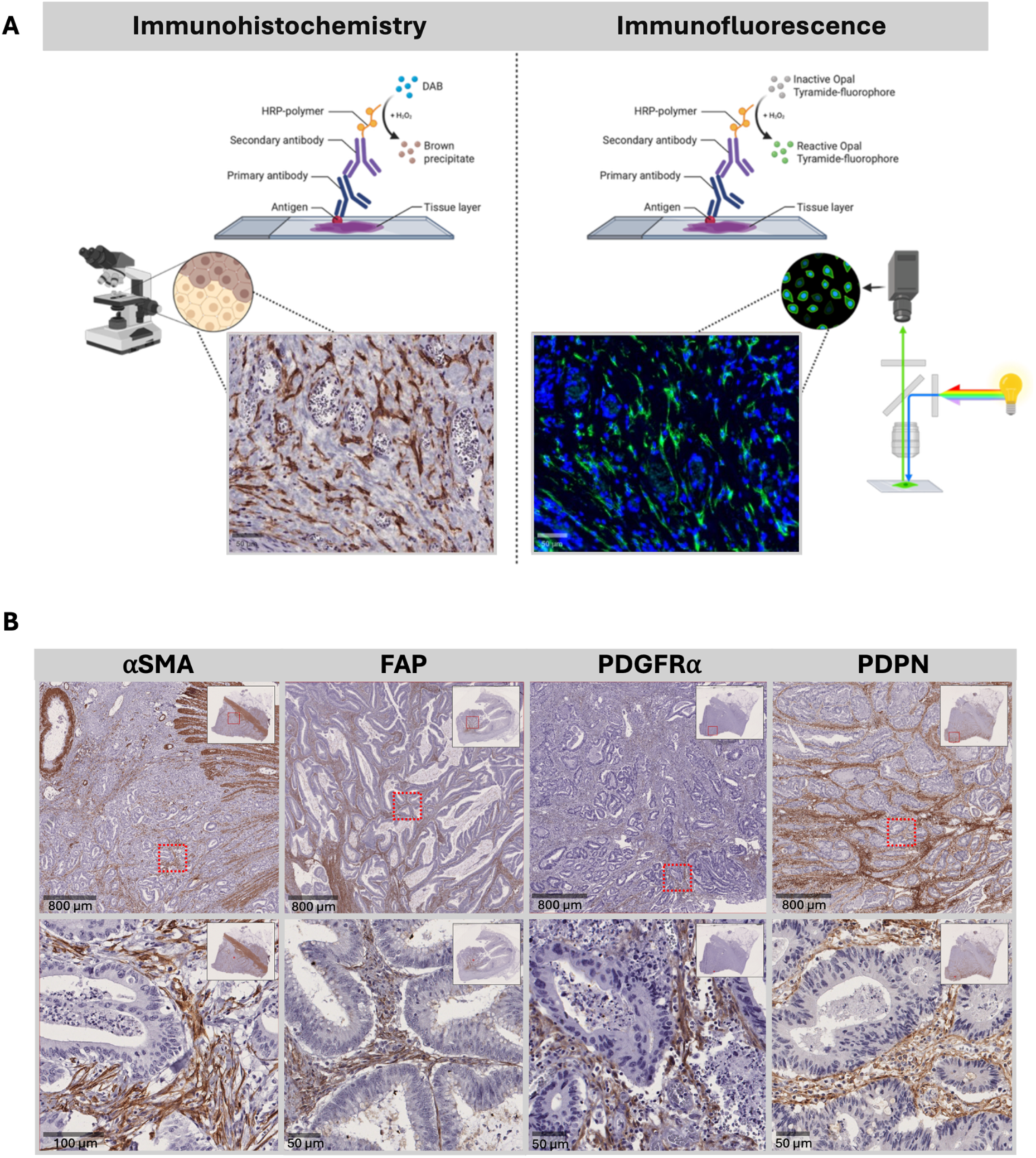
Immunohistochemistry optimisation in colorectal carcinoma. A schematic of the antibody complex established in an IHC vs sIF staining procedure (A). Example images of colorectal carcinoma control tissue stained with the optimal IHC conditions for each CAF antigen of interest. Red boxes highlight regions displayed at higher resolution in the lower panel (B).

Whilst modest relative to the high plex platforms described above, the 6-plex panel described in this report facilitates a focussed approach to profiling CAFs and allows researchers to assess their spatial relationship with CD8+ T cells and panCK+ cancer cells. Below we have outlined the experiments that are required to develop a reliable multiplex panel and highlight the importance of careful design and rigorous optimisation. In brief, the steps include optimisation of IHC conditions, testing of conditions by single-plex immunofluorescence (sIF), confirmation of antibody heat inactivation, optimisation of antibody position within a multiplex cycle, robust selection of antibody to Opal pairs and finally, multiplex immunofluorescence (mIF) panel testing.

## 2. Materials and Methods

### 2.1. Samples

Protocol optimisation was conducted on formalin-fixed paraffin-embedded (FFPE) colorectal carcinoma control tissue. Samples from the RADIO trial (ISRCTN43698103) represent untreated muscle-invasive bladder cancer (MIBC) tissue collected during diagnostic transurethral resections of bladder tumours. All patients provided informed consent for tissue collection and research on RADIO samples received ethical approval (REC: 23/NE/0212). All wet lab work described was completed in the Integrated Pathology Unit (IPU) at the Royal Marsden NHS Hospital Trust and the Institute of Cancer Research, London. Staining was reviewed by pathologist Dr. Tatiany Silveira (TS).

### 2.2. Tissue preparation

FFPE tissue was cut in 3 μm sections. Prior to IHC, sIF or mIF slides were baked overnight at 50°C, followed by 1 hour at 62°C on the day of staining. For single plex protocols, slides were manually de-waxed in xylene and sequentially rehydrated in 100%, 95%, 70% and 50% ethanol. For multiplex staining the dewaxing and rehydration steps were completed using the Leica BOND 3.0 RX/RXm automated immunostainer (Leica Microsystems, Milton Keynes, UK).

### 2.3. Chromogenic immunohistochemistry

Single plex IHC was performed on FFPE tissue sections for each of the markers individually to ascertain the primary antibody concentration and the optimal conditions for heat-induced epitope retrieval (HIER). After baking, deparaffinization and rehydration, slides were subjected to microwave HIER in citrate buffer solution at pH 6.0, or EDTA at pH 9.0 (Invitrogen low pH 00-4955-58 / high pH 00-4956-58), first for 1 minute and 30 seconds at 1000 watts and then 15 minutes at 100 watts. This was followed by a 10-minute peroxidase block with 3% hydrogen peroxide (Sigma Aldrich, 88597). Slides were washed in Tris-Buffered Saline, 0.1% Tween ® 20 (Sigma, P9416) (TBST) and subjected to 1-hour protein block [serum-free protein block (Dako, X0909)]. To establish the optimal antibody dilution, a serial dilution of each primary antibody was prepared in Primary Antibody Diluent (Leica Biosystems, AR9352) and applied for 1-hour at room temperature. Slides were washed with TBST and incubated with a mouse or rabbit specific secondary HRP conjugated secondary antibody (SignalStain® Boost IHC Detection Reagent HRP, Cell Signaling, Rabbit 8114S/ Mouse 8125S) according to the species of the primary antibody Fc region. Slides were washed with TBST and DAB solution (Novolink Max DAB polymer, Leica Biosystems, RE7270) was applied in a dark chamber. Finally, slides were rinsed in deionized water, counterstained with hematoxylin (Shandon Harris, REF6765004), dehydrated and mounted with Cytoseal 60 (Thermo Scientific, 60 8310-16).

### 2.4. Immunohistochemistry evaluation by pixel classification

To evaluate antigen stability after multiple rounds of HIER, a pixel classifier was trained in QuPath (version 0.5.1) to detect positive DAB staining for each antigen of interest (json files for each pixel classifier are provided in supplementary files). Three regions were selected in the MIBC control tissue stained for each antigen. In each region annotations were added to provide examples of positive and negative staining. The performance of the classifier was considered whilst annotations were added to assess the accuracy and make modifications as required.

### 2.5. Single plex immunofluorescence

To detect the antigens of interest by immunofluorescence, the Opal-TSA system was adopted using the Opal Polaris 7-Color Automation IHC Kit (Quanterix, NEL871001KT). Single plex immunofluorescence staining was performed manually following a similar procedure to the IHC protocol (Figure 1A) with adjustments to the reagents and incubation times as per the Opal kit guidelines (Supplementary Table 1). In brief, slides were baked, deparaffinized, rehydrated and subjected to microwave HIER. Slides underwent a 10-minute peroxidase block and were washed with TBST, followed by a 10-minute protein block. Slides were incubated with the primary antibody for 30 minutes at room temperature then washed with TBST. The secondary antibody was applied for 10 minutes then washed with TBST. In rare cases where detection was less efficacious in the Opal system, a secondary polymer (Leica Biosystems, RE7200-K) was utilised, and the primary antibody dilution adjusted. The diluted Opal reagent was subsequently applied for 10 minutes. For single plex optimisation, all antibodies were paired with Opal 520 to assess the IHC-IF concordance. The slides were washed in TBST and subjected to an additional round of HIER prior to application of 4′,6-diamidino-2-phenylindole (DAPI) nuclear stain and mounting with water-based mounting medium (Shandon, Immu-MountTM, 9990402). The resulting images were compared to matched IHC stained slides and visually assessed by a pathologist (TS).

### 2.6. Automated multiplex immunofluorescence

All multiplex staining protocols were completed using the Leica BOND 3.0 RX/RXm automated immunostainer. A bespoke protocol utilising ancillary reagents, was created to facilitate the inclusion of multiple antibody and Opal reagents. As with previous protocols, the slides were baked in advance then loaded into a BOND rack for the staining. Briefly the protocol is programmed to complete the following steps: 1. tissue deparaffinization and rehydration, 2. HIER, 3. peroxidase and protein blocking, 4. primary antibody staining, 5. secondary antibody staining, 6. application of Opal and all appropriate washes. Steps 2-6 are repeated for a further 5 cycles. The slides are then manually washed and counterstained with DAPI for 5 minutes at room temperature before mounting with water-based mounting medium.

To conduct the fluorescence minus one experiment, seven control slides were stained with the multiplex panel using the Bond autostainer. The first slide was stained with the full multiplex panel, for each subsequent slide the Bond protocol was adapted to replace one primary antibody with a Bond wash step to generate a secondary only control (Supplementary Table 3).

### 2.7. Image acquisition

IHC stained slides were scanned in brightfield mode with the Hamamatsu Nanozoomer at 20X. Immunofluorescence-stained slides were scanned on the PhenoImager (formerly Vectra Polaris version 1.0.13; Quanterix) at 20X.

Following image acquisition, mIF images were passed through the spectral unmixing software InForm (version 2.6.0; Quanterix). Post InForm, the output component image tiles were stitched to recreate the whole slide image using a stitching script generated by Pete Bankhead for QuPath [17].

### 2.8. Immunofluorescence evaluation by pixel classification

To evaluate staining in the fluorescence minus one experiment, QuPath was used to generate a single channel threshold for each antigen to detect the percentage of positive pixels within a selected region. Supplementary Table 4 documents the threshold value set for each antigen, where pixels above the threshold value were classed as positive.

## 3. Results

### 3.1. Optimising immunohistochemistry conditions for CAF markers of interest

A thorough literature review was conducted across multiple tumour types to identify frequently cited panCAF markers, and to facilitate identification of CAF subtypes and states at the level of both protein and RNA. Selected antigens included αSMA, CD90, fibroblast activation protein (FAP), platelet derived growth factor receptor alpha (PDGFRα) and beta (PDGFRβ), podoplanin (PDPN), Regulator of G-protein Signaling 5 (RGS5), S100A4 and Vimentin. Where possible, selection of antibodies and specific clones was guided by published evidence of strong and specific staining. Of note, the antibodies we acquired were unconjugated and therefore suitable for downstream use in the Opal-TSA system.

Prior to multiplexing, the performance of antibodies by IHC was established. Doing so generates a reliable reference point to ensure the immunofluorescence staining patterns are as expected and not an artefact or due to autofluorescence. Each CAF antibody was assessed by IHC to identify the optimal staining conditions for protein detection in stromal regions and presence on elongated cells with spindle-shaped fibroblast morphology was confirmed (Figure 1B) (Table 1). Initial experiments were conducted in control tissue containing colorectal carcinoma due to the abundant stromal content and reported expression of several of the CAF markers of interest [18]. IHC and mIF conditions for CD8 and pan-cytokeratin (panCK) were previously optimised (data not shown). The performance of CAF antibodies was also validated in five control MIBC cases (Figure 2, Supplementary figure 1) to ensure that staining conditions were effective across heterogenous bladder samples and to explore which antigens were more likely to be expressed in unique areas or co-expressed with other antigens in the panel.

**Figure 2:**
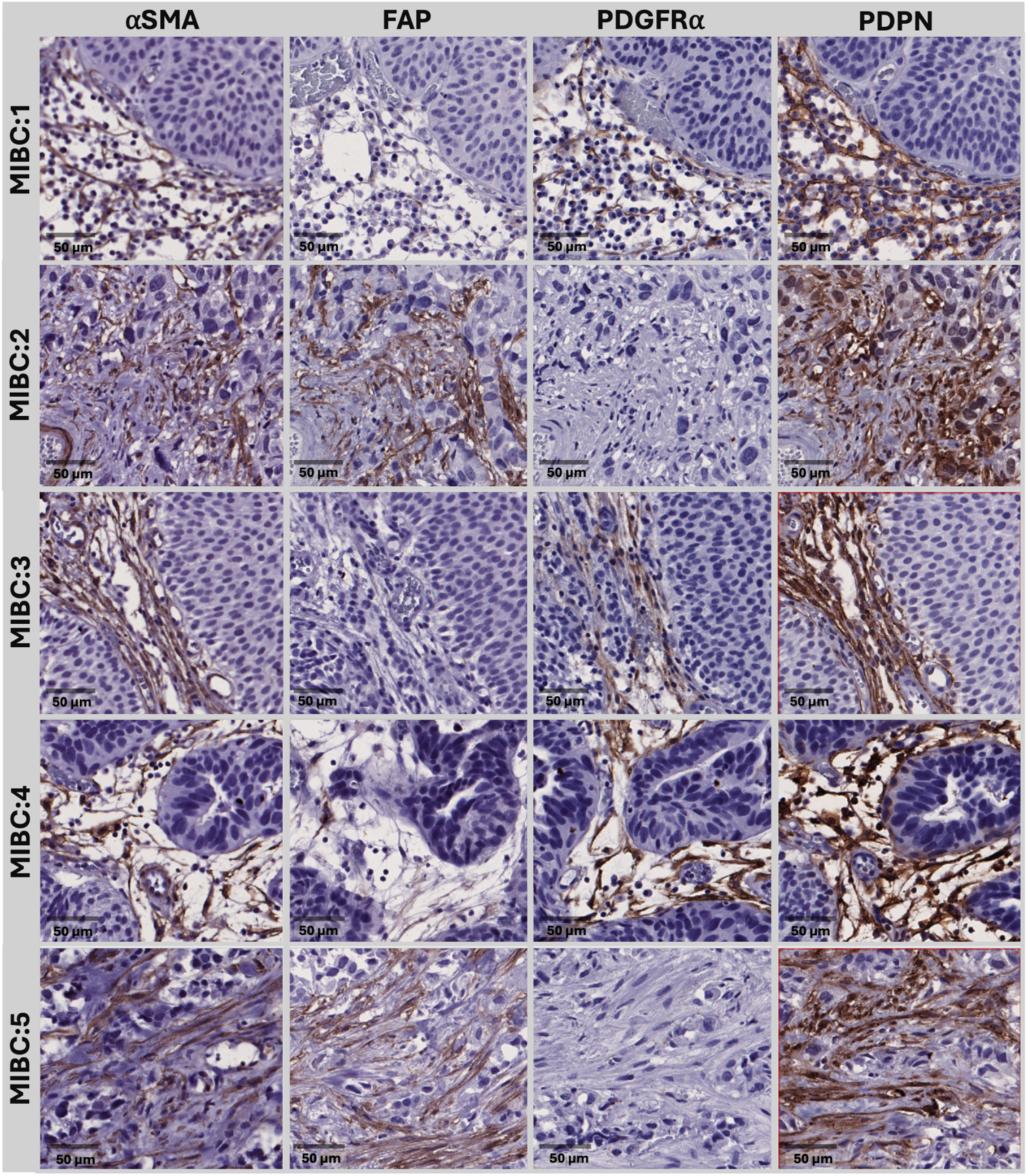
Validation of immunohistochemistry conditions in muscle-invasive bladder cancer. To validate the optimal IHC conditions, five MIBC samples were stained with the CAF markers of interest. Each row represents a randomly selected region of tissue from a unique MIBC control, each column highlights which antigen was stained for, positive staining is shown in brown (scale bar = 50 µM).

**Table 1:**
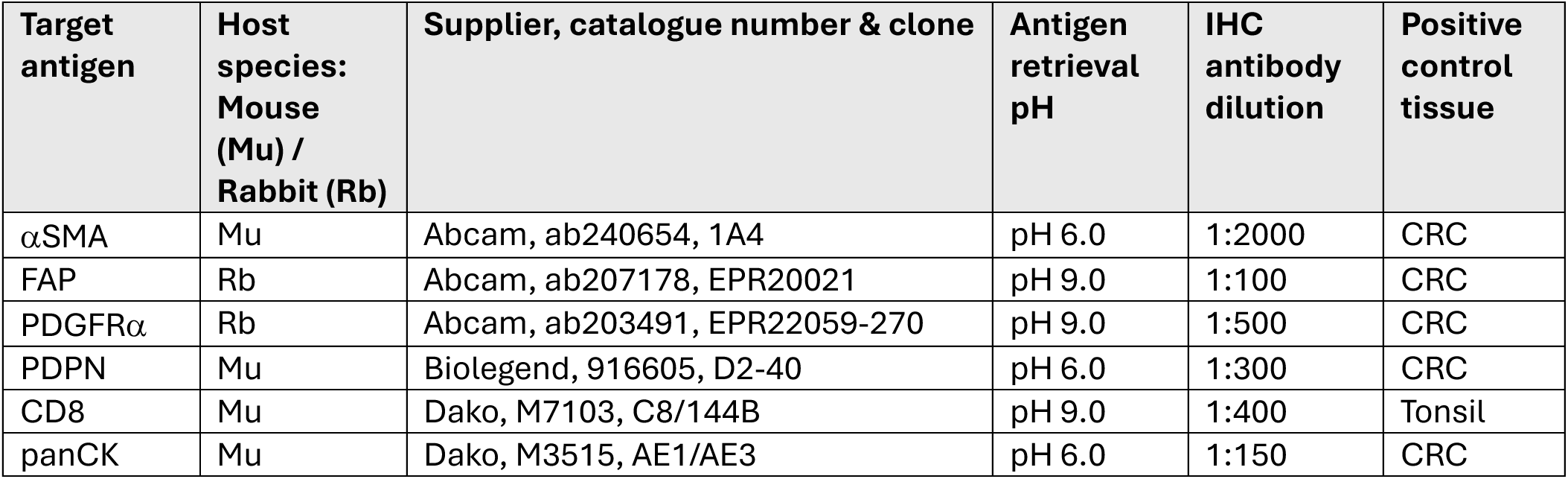
Optimal IHC conditions for antigens of interest as established in control tissue.

| Target antigen | Host species:<br>Mouse (Mu) /<br>Rabbit (Rb) | Supplier, catalogue number & clone | Antigen retrieval pH | IHC antibody dilution | Positive control tissue |
| --- | --- | --- | --- | --- | --- |
| αSMA | Mu | Abcam, ab240654, 1A4 | pH 6.0 | 1:2000 | CRC |
| FAP | Rb | Abcam, ab207178, EPR20021 | pH 9.0 | 1:100 | CRC |
| PDGFRα | Rb | Abcam, ab203491, EPR22059-270 | pH 9.0 | 1:500 | CRC |
| PDPN | Mu | Biolegend, 916605, D2-40 | pH 6.0 | 1:300 | CRC |
| CD8 | Mu | Dako, M7103, C8/144B | pH 9.0 | 1:400 | Tonsil |
| panCK | Mu | Dako, M3515, AE1/AE3 | pH 6.0 | 1:150 | CRC |

CAF markers identified by single cell RNA sequencing warrant consideration and IHC evaluation enables confirmation that proteomic expression is consistent with the expected cell type. Regulator of G Protein Signaling 5 (RGS5) has been previously reported as a potential marker of myCAFs in MIBC using scRNASeq [9], however, we observed considerable staining of both tumour and stromal cells using RGS5. We also observed tumour cell staining using antibodies against S100A4 and pan-stromal staining with Vimentin (Supplementary Figure 1). Whilst the difference between tumour and stromal cells expressing antigens could likely be resolved through regional demarcation and image analysis techniques, when limited to a few markers to identify CAFs on the multiplex panel, pragmatic considerations meant these less specific antigens were excluded.

In contrast, PDPN staining was abundant in the stromal regions in all MIBC controls. Likewise, staining for αSMA could be detected in the stroma of all MIBC controls, but to varying degrees. The stromal expression of FAP and PDGFRα appeared to be mutually exclusive, potentially highlighting the presence of different CAF subtypes with heterogenous spatial arrangements in the tumour microenvironment. Based on the observed staining patterns, the multiplex panel was refined from a list of 10 possible markers to include four fibroblast markers: αSMA, FAP, PDPN, and PDGFRα in addition to the cytotoxic T cell marker CD8 and the tumour cell marker panCK. In combination, the four selected CAF markers were considered sufficient to identify the majority of CAFs in the tumour microenvironment, with the potential to further identify distinct CAF subpopulations. This series of IHC experiments affirms the need to consider multiple CAF markers to capture heterogeneity within the stroma. In addition, it highlights the need to carefully select the control tissue and to consider multiple regions of interest if heterogeneity within the cell population of interest is expected.

### 3.2. Antigen retrieval conditions to determine the antibody position

During the multiplex process, antigens are stained sequentially and are subject to several rounds of HIER. To better understand the stability of each antigen of interest and thus predict the optimal position in the multiplex protocol, IHC was performed with 3 control slides at the optimised conditions for the primary antibody, with the following modifications in HIER: one slide to evaluate antibody position 1, submitted to 1 cycle of HIER, one slide to evaluate antibody position 3, submitted to 3 cycles of HIER and one slide to evaluate antibody position 5, submitted to 5 cycles of HIER.

Antigen stability was assessed in QuPath by quantifying the percentage of positive pixels in three comparable regions (Figure 3). Regions included in the quantification are shown in Supplementary Figures 2-5. The detection of αSMA was deemed stable and had minimal change in the percentage of positive pixels under all HIER conditions, therefore allowing αSMA to be placed in any position in the panel. In contrast, FAP could only withstand one round of HIER, additional rounds of HIER resulted in a near complete depletion of detectable antigen. To avoid epitope denaturation, FAP needed to be placed in position one in the multiplex panel. Whilst there was some depletion of stromal PDPN staining from HIER 1 to 3, the general staining pattern was maintained with minimal difference between HIER 3 and 5. PDGFRα was the only antigen where specific stromal detection improved with more rounds of antigen retrieval. To note, the control tissue used for this study contained an isolated area with limited PDGFRα expression, therefore selection of three distinct areas to quantify positive pixels was limited. Supplementary Table 2 summarises the available positions for each antigen of interest. This experiment demonstrates the differences in antigen stability and highlights the importance of considering how HIER can impact the order of staining in a multiplex panel.

**Figure 3:**
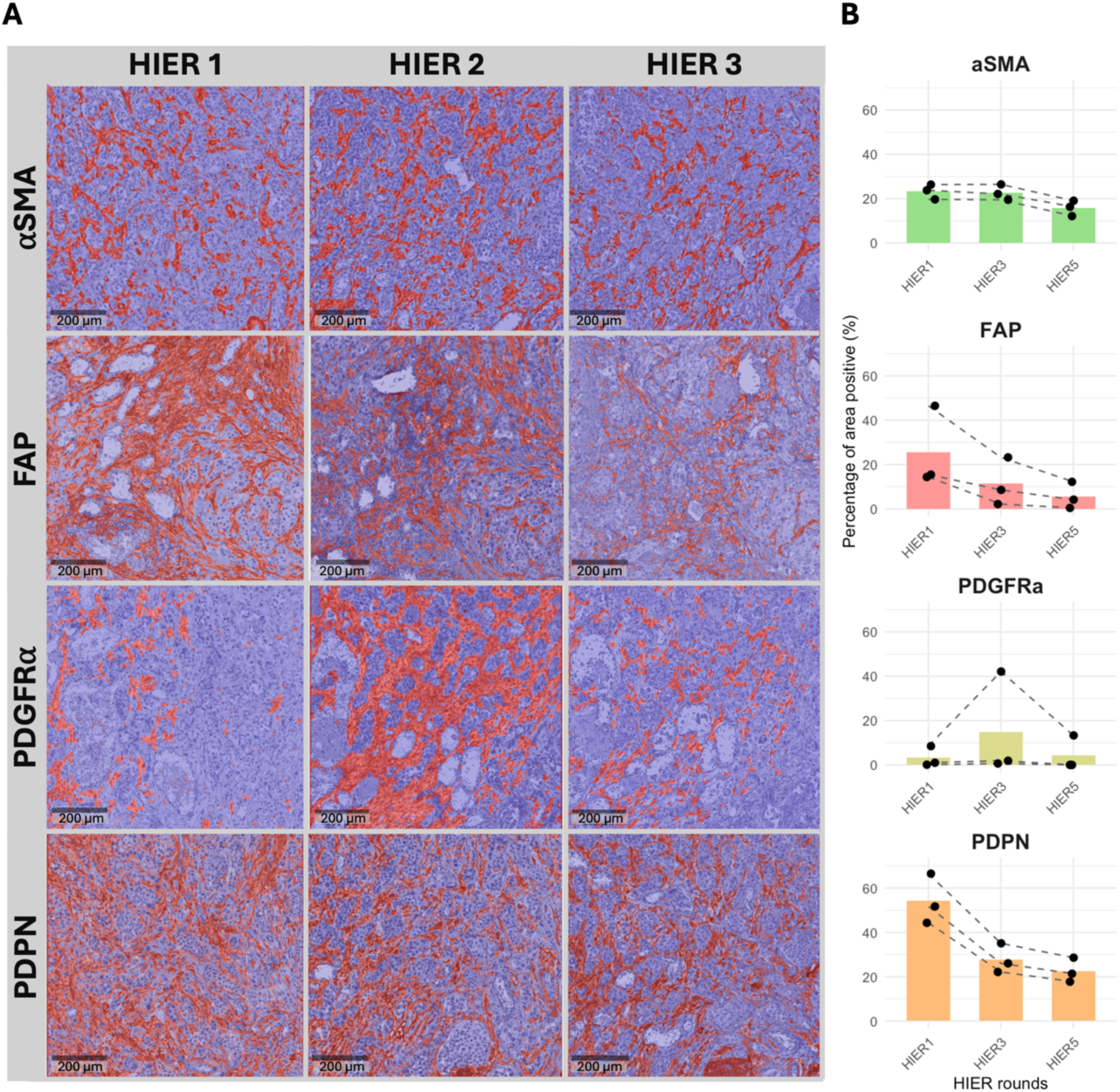
Assessment of antigen tolerance to heat-inactivation/ epitope retrieval. Example images show MIBC tissue stained with the optimal conditions for each antigen of interest following 1, 3 and 5 rounds of HIER. IHC-stained images are overlayed with the QuPath positive pixel detection of DAB staining shown in red (A). The percentage of positive pixels was quantified in three 1000x1000 pixel regions and compared following each round of HIER (B), (scale bar = 200 µM).

### 3.3. Assessing the efficacy of heat inactivation and epitope retrieval

The HIER step is multifunctional; it exposes the antigens of interest for optimal detection and crucially, for mIF, the HIER clears the antibody complex from the previous staining round preparing the tissue for subsequent staining cycles. If ineffective and an antibody remains bound to the tissue, it can be detected by multiple Opals leading to confusing results (Figure 4B).

**Figure 4:**
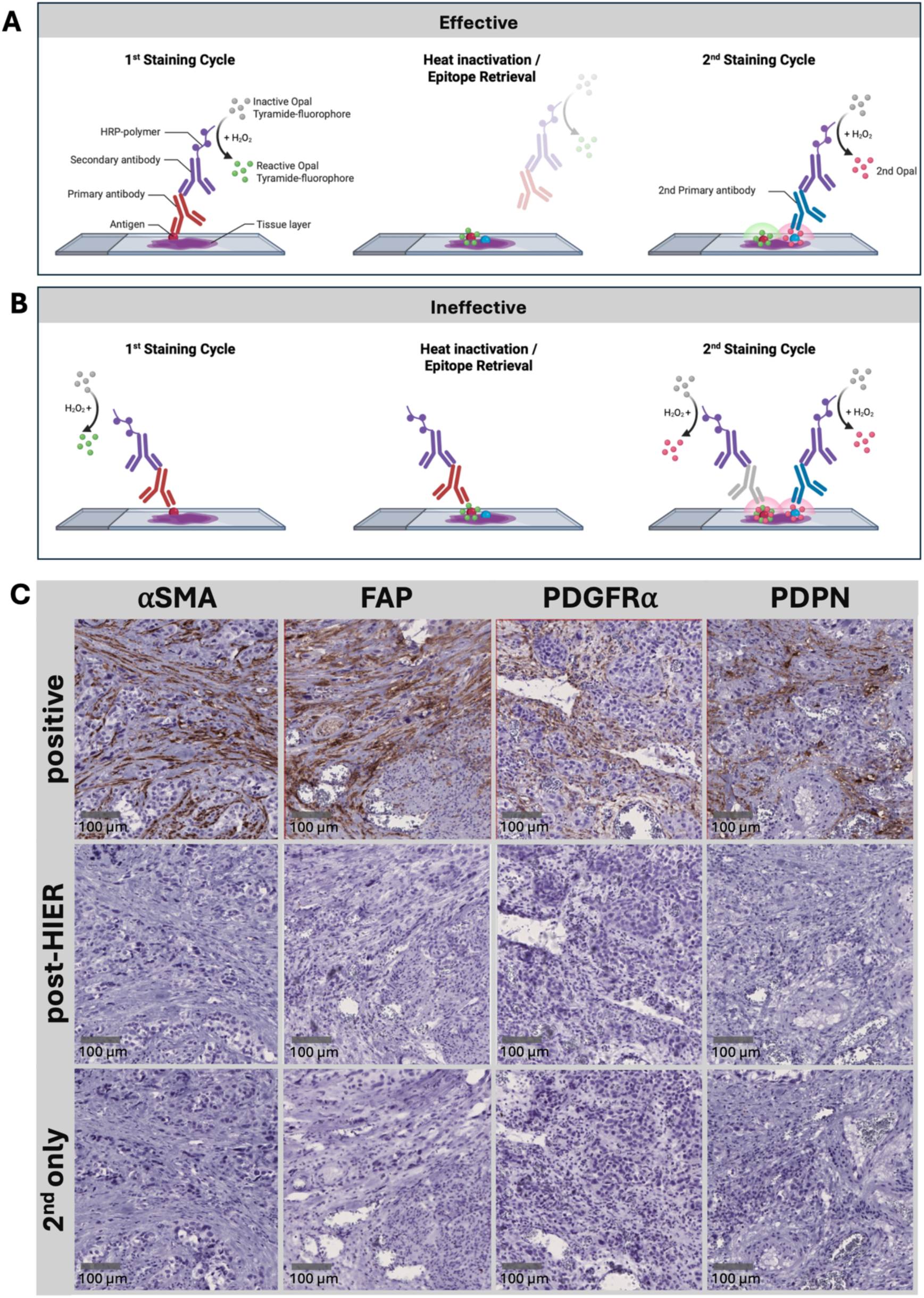
Assessment of heat-inactivation/ epitope retrieval efficiency. Schematic showing the treatment with HIER as an essential step in mIF to remove the antibody complex prior to the next staining cycle (A). Inefficient HIER leads to residual positive staining of the first antibody within the second staining cycle (B). Example images show MIBC tissue stained with the optimal conditions for αSMA, FAP, PDGFRα, and PDPN, either without HIER treatment (*positive control*), after the HIER protocol was applied (*post HIER*) or with a secondary only control (*2nd only control*) (C). With effective HIER the image should appear the same as the secondary only control, (scale bar = 100 µM).

To ensure that the HIER step effectively cleared the antibody from the tissue, the following conditions were evaluated for each antibody on concomitant slides: one positive control slide stained with standard primary and secondary antibodies, one negative secondary only control slide and one test slide subjected to primary and secondary staining, followed by a 20-minute HIER and subsequent DAB detection. If 20 minutes of HIER is sufficient to clear the antibody complex, the test condition should mirror the negative control. If ineffective, the HIER time should be extended. Qualitative assessment of the images in Figure 4C confirmed that that 20 minutes HIER was sufficient to clear the antibody complexes and prevent subsequent detection. This confirmation of effective removal of antibodies is essential to ensure robust staining patterns are produced with the multiplex panel.

### 3.4 Validation of staining conditions for single plex immunofluorescence

Building on the optimal IHC conditions, sIF staining was optimised to compensate for differences between the IHC and IF protocols and to account for additional signal amplification that is typically observed with the Opal system reagents. Optimal conditions are shown in Figure 5. The staining patterns detected by sIF reflected those seen by IHC in the same control tissue. To note, PDPN failed to be detected by sIF with the use of the Quanterix secondary antibody (Supplementary Figure 6). This was rectified with the Leica post primary and polymer system which restored detection and resulted in comparable staining patterns to the IHC control. Due to the amplification of signal with the polymer, the antibody was further diluted to 1:800 for further experiments. Reassuringly, the polymer only control did not result in any non-specific immunofluorescence detection. This experiment highlights the value of the reference IHC image to validate the immunofluorescence staining patterns.

**Figure 5:**
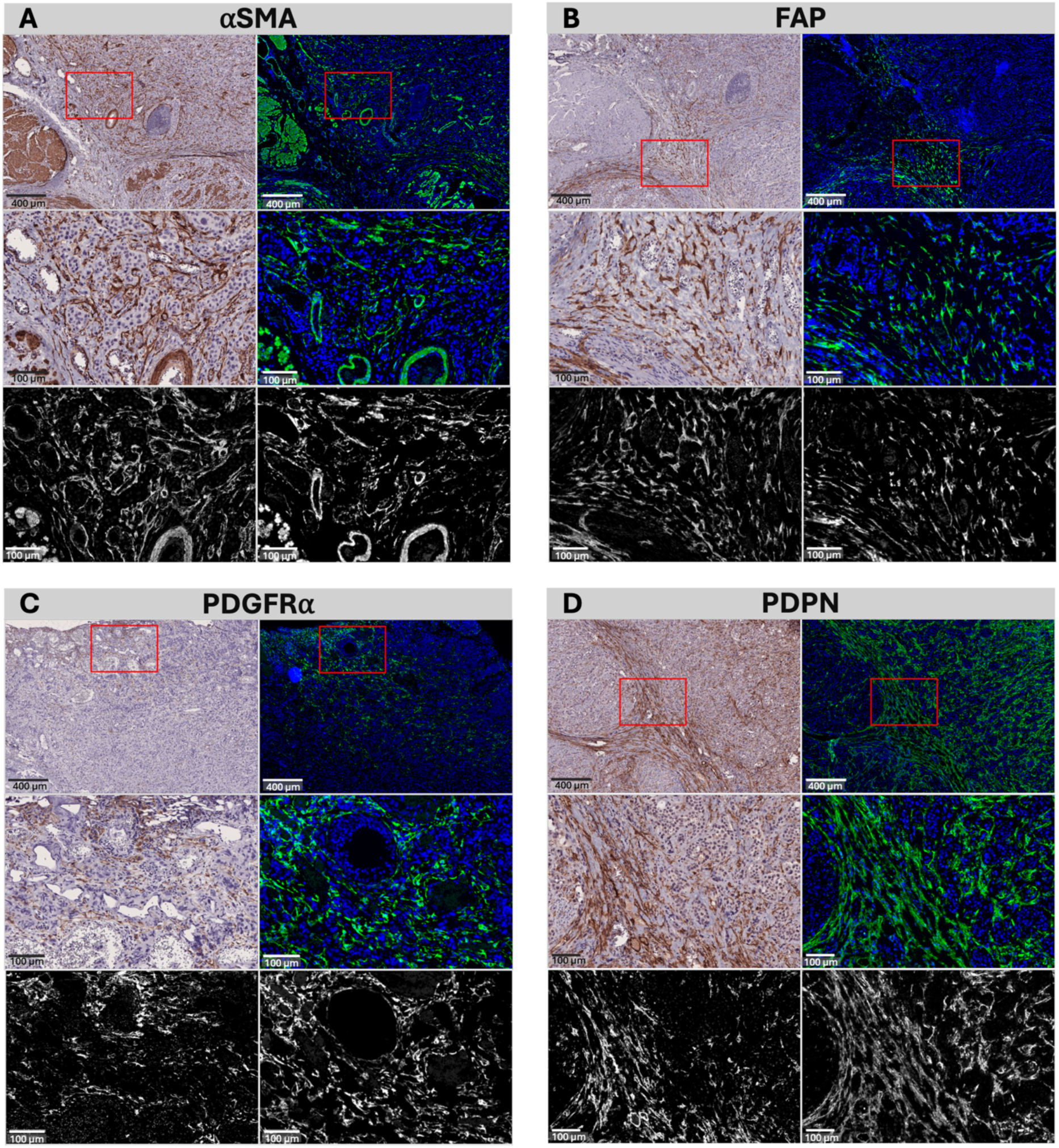
Validation of staining conditions by single plex immunofluorescence. Example images of MIBC tissue stained by IHC (left) or sIF (right) for each CAF antigen of interest (A-D). In each panel a low-resolution overview is shown on the top row, (scale bar = 400 µM). Red boxes highlight the zoomed insert shown in the middle row, (scale bar = 100 µM). A duplicate of the insert is displayed in grayscale in the bottom row to demonstrate the similarity in staining patterns irrespective of the IHC and sIF display colours.

### 3.5 Testing of the final multiplex panel design

The final multiplex panel was designed taking into consideration the antibody concentration, pH, antigen stability and performance in the Opal system as outlined in the results above. In addition, antigen expression and cellular localisation was evaluated to inform suitable antibody-Opal pairings (Table 2), where antigens with low expression were paired with the brightest Opals. Example images of the multiplex panel test are displayed in Figure 6A. Here, two spatially-distinct regions highlight the heterogeneity of CAF marker expression within the same tissue. Application of the panel to the five MIBC control samples further highlighted the inter-sample heterogeneity that could be detected with the panel (Figure 6B).

**Figure 6:**
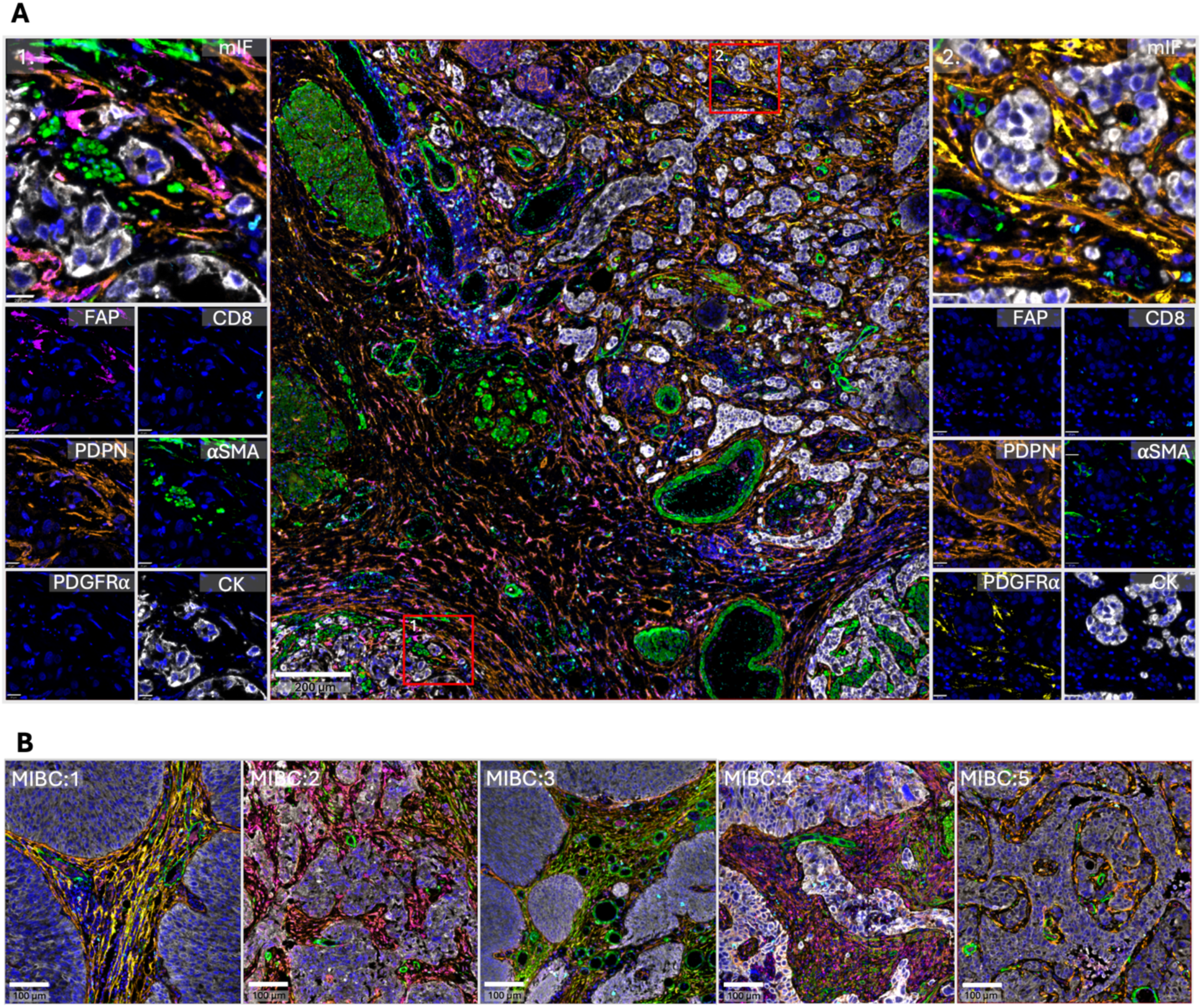
Assessment of the final multiplex panel design. The central image provides an example of a large region of MIBC stained with the optimised CAF mIF panel (scale bar = 200 µM). Red squares highlight two insert regions with heterogenous staining patterns within a single tissue. Insert 1 depicts a region with increased expression of FAP, whereas insert 2 displays a region with enrichment of PDGFRα (scale bar = 20 µM). Below each insert the same region is shown with each individual antigen. To highlight inter-sample heterogeneity, five MIBC control samples were stained with the CAF mIF, each tile represents a randomly selected region of tissue from each sample (scale bar = 100 µM) (B).

**Table 2:**
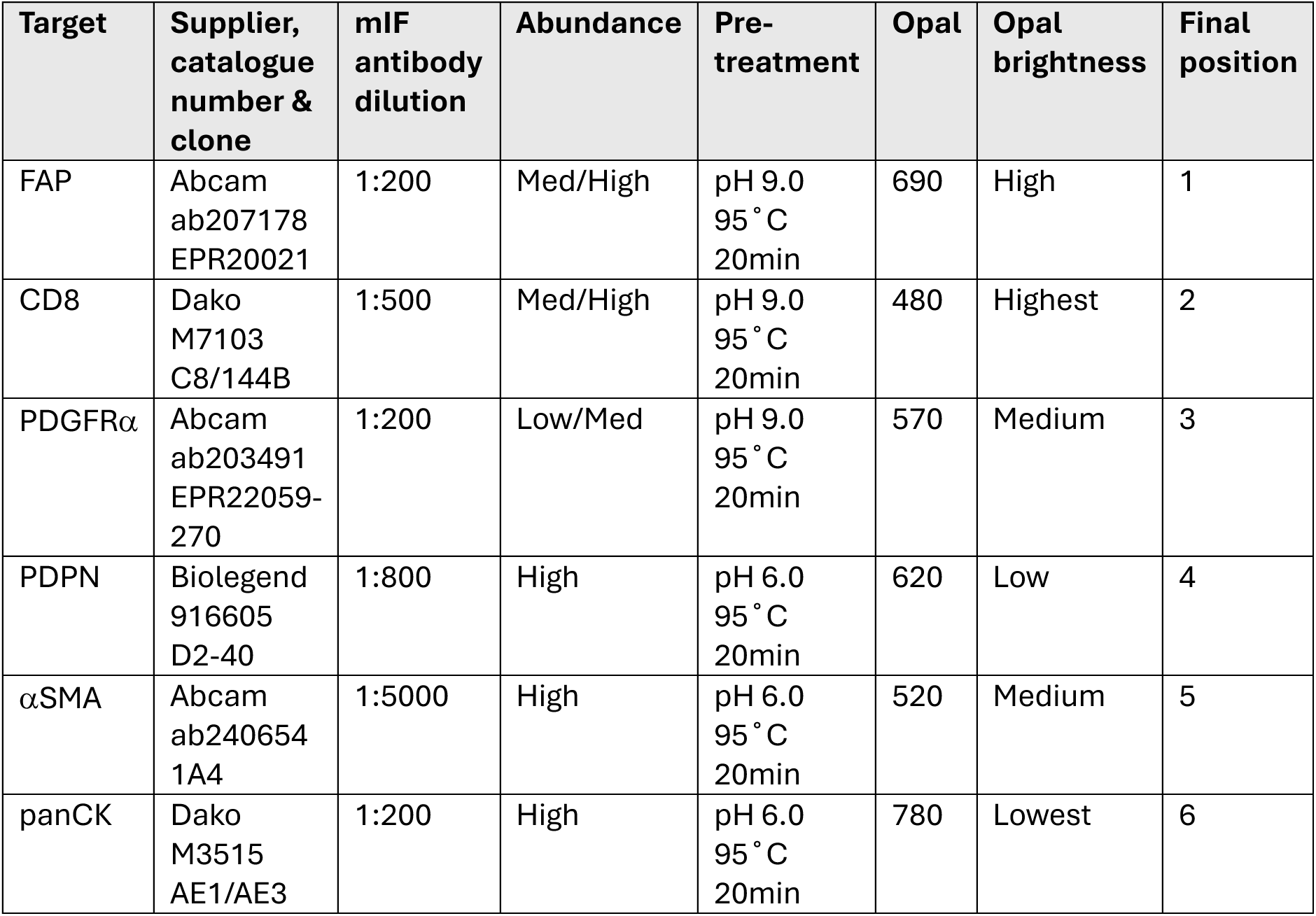
A summary of the fully optimised multiplex panel antibody conditions and Opal pairings.

To further evaluate the efficacy of the HIER step in the multiplex setting, the full panel was tested in a fluorescence minus one (FMO) experiment with a series of secondary only controls. Detection of an Opal in the control position suggests that an antibody from a previous step is still bound to the tissue and the HIER step is not fully efficacious in clearing the antibody. Qualitative and quantitative assessment confirmed the absence of antigen detection in the secondary only positions, confirming that the antibody complex was effectively cleared during each HIER step in the staining cycle (Supplementary Figures 7,8,9). To note, due to the intra-sample heterogeneity within the control tissue used for the FMO study two regions were selected for assessment. Like the IHC-based HIER tests, the FMO experiment is critical for building confidence in the final staining patterns and ensures that antibodies are effectively cleared from the tissue, preventing detection in subsequent staining cycles.

## 4. Discussion

Following careful optimisation, we have demonstrated that the 6-marker multiplex immunofluorescence panel defined in this report can be applied to routinely-available FFPE tissue to reliably and reproducibly generate high quality images to accurately profile CAFs in the tumour microenvironment. Whilst we acknowledge that the inclusion of only four CAF markers on the panel restricts the level of biological complexity that can be observed, this approach allowed us to characterise a variety of CAF phenotypes that express different combinations of the four CAF markers. This level of diversity could not be observed with a single marker and would be difficult to detect by IHC staining on sequential sections. In addition, the high-throughput nature of this multiplex approach means that large whole slide images can be generated for many samples, enabling comprehensive evaluation of both intratumoral and intertumoral heterogeneity. On average we were able to stain and scan 30 whole slide images in 48 hours.

In this report, panel testing was primarily conducted in MIBC tissue, however, the selected markers have been frequently cited in CAF literature across different tumour types, we anticipate that the panel is suitable to study CAFs in other tumour types. In addition, the experiments we have outlined provide guidance for researchers planning to develop their own bespoke panel.

A major advantage of mIF is the preservation of tissue architecture to understand the arrangement of the tumour microenvironment at single cell resolution. Access to the spatial coordinates of cells provides the opportunity to explore the spatial relationships between cell types of interest. Many spatial analysis approaches build upon the principles of topology, geometry and ecology to map patterns within cell populations to quantify the complexity and organisation of tissues. Popular techniques include nearest neighbour analysis and the identification of recurrent cellular neighbourhoods to elucidate spatially-distinct clusters of cells. These can be powerful approaches to understand local cellular interactions and associated paracrine signalling in the TME.

Single cell RNA sequencing has been instrumental in defining CAF subtypes and function [19]. This approach provides quantitative data for thousands of genes and is often used to infer cellular function, however, gene activity does not perfectly correlate with expression of functional proteins. In contrast, mIF facilitates quantification of actual protein expression and is more likely to reflect a true phenotypic state. There are benefits to both transcriptomic and proteomic approaches and the insights from each are complimentary and not interchangeable. There has been recent exponential growth and interest in spatial transcriptomics platforms leading to the generation of single cell spatial datasets across multiple tumour types [10]. As a result, CAFs can be profiled with increasing detail leading to definition of many more CAF subtypes beyond iCAFs and myCAFs, such as CD74+ antigen-presenting CAFs [2, 20] or perineural neuronal-CAFs [21]. These rapidly-emerging technologies often combine multiplex immunofluorescence with transcriptomics, for example Bruker CosMX, Lunaphore COMET, Leica CellDIVE and MACsima to maximise proteomic and transcriptomic readouts with spatial single cell resolution. Such techniques rely on high quality, well optimised mIF and some of the key considerations outlined in this report are relevant to their robust use.

The generation of huge, multilayered, single cell spatial datasets provides exceptional opportunities for biologically complex analyses, however, increasing complexity brings new challenges, particularly relating to computational requirements of image processing, data analysis and storage. There is increasing availability of open-source software for analysis such as Qupath [17] and to streamline spatial analysis, including generation of resources such as the Steinbock toolkit [22], TRACERx-PHLEX [23] and MuSpAn [24]. With increasing adoption of artificial intelligence (AI), it seems inevitable that advances in future image analysis techniques will include AI approaches. Deep-learning models, such as NaroNet, have been developed with the ability to learn cell phenotypes and the arrangement of cellular neighbourhoods in multiplexed images [25]. Similarly, by using information from multiplex images, generative AI offers exciting opportunities to predict the expression of biomarkers and generate highly-plexed images from simple and affordable Hematoxylin and Eosin-stained images [26]. Such tools typically require large training datasets but, in the future, may be able to assist with the generation and interpretation of multiplexed images, making the technique more widely accessible, including to the clinic.

## 5. Conclusions

Through careful panel design and optimisation, we have shown that multiplex immunofluorescence is well suited to the study of heterogenous cell types, such as CAFs, where multiple markers are required to capture diversity within the population. Using a combination of αSMA, FAP, PDPN, and PDGFRα we have shown that multiple CAF phenotypes can be detected with spatially distinct single cell resolution and have outlined comprehensive details of the CAF panel so that it can be utilised by other researchers. By combining these CAF markers with CD8 and panCK, spatial analysis approaches can assess how their colocalization with distinct CAF subtypes may contribute to pro-tumorigenic functions of CAFs and immune exclusion. This provides insights that are relevant for future CAF-targeting strategies. Finally, for researchers planning to generate their own bespoke mIF panel, this report outlines the multiple steps of the optimisation protocol. It emphasises the need for critical assessment of images with cross referencing to a gold-standard IHC and specific control experiments to generate reliable multiplex images for large-scale analysis.

## Supporting information

Supplementary json files

## 6. Author contributions

AB, TS, TL and AW contributed to conceptualisation. AB conducted staining, data collection and image analysis. TS conducted pathological assessment of images and experimental supervision. AB, TS and AW contributed to manuscript writing and editing. All authors reviewed and approved the final manuscript.

## 7. Acknowledgements

This manuscript represents independent research supported by the National Institute for Health Research (NIHR) Biomedical Research Centre at The Royal Marsden NHS Foundation Trust and the Institute of Cancer Research, London. The views expressed are those of the author(s) and not necessarily those of the NIHR or the Department of Health and Social Care. The authors acknowledge funding from Cancer Research UK RadNet at The Institute of Cancer Research and The Royal Marsden Hospital and City of London RadNet.

## Abbreviations

(CAFs): Cancer-associated fibroblasts
(myCAFs): myofibroblasts-like CAFs
(iCAFs): inflammatory CAFs
(αSMA): alpha smooth muscle actin
(IHC): immunohistochemistry
(TSA): tyramide signal amplification
(sIF): single-plex immunofluorescence
(mIF): multiplex immunofluorescence
(FFPE): formalin-fixed paraffin-embedded
(MIBC): muscle-invasive bladder cancer
(HIER): heat-induced epitope retrieval
(TBST): Tris-Buffered Saline, 0.1% Tween ® 20
(DAPI): 4′,6-diamidino-2-phenylindole
(FAP): fibroblast activation protein
(PDGFRα): platelet-derived growth factor receptor alpha
(PDGFRβ): platelet-derived growth factor receptor beta
(PDPN): podoplanin
(RGS5): Regulator of G-protein Signaling

## References

1. Muhl, L., et al., Single-cell analysis uncovers fibroblast heterogeneity and criteria for fibroblast and mural cell identification and discrimination. Nat Commun, 2020. 11(1): p. 3953.

2. Song, J., et al., Antigen-presenting cancer associated fibroblasts enhance antitumor immunity and predict immunotherapy response. Nature Communications, 2025. 16(1).

3. Friedman, G., et al., Cancer-associated fibroblast compositions change with breast cancer progression linking the ratio of S100A4(+) and PDPN(+) CAFs to clinical outcome. Nat Cancer, 2020. 1(7): p. 692–708.

4. Bhattacharjee, S., et al., Tumor restriction by type I collagen opposes tumor-promoting effects of cancer-associated fibroblasts. Journal of Clinical Investigation, 2021. 131(11).

5. Foster, D.S., et al., Multiomic analysis reveals conservation of cancer-associated fibroblast phenotypes across species and tissue of origin. Cancer Cell, 2022.

6. Öhlund, D., et al., Distinct populations of inflammatory fibroblasts and myofibroblasts in pancreatic cancer. The Journal of experimental medicine, 2017. 214(3): p. 579–596.

7. Costa, A., et al., Fibroblast Heterogeneity and Immunosuppressive Environment in Human Breast Cancer. Cancer Cell, 2018. 33(3): p. 463–479.e10.

8. Obradovic, A., et al., Immunostimulatory Cancer-Associated Fibroblast Subpopulations Can Predict Immunotherapy Response in Head and Neck Cancer. Clin Cancer Res, 2022. 28(10): p. 2094–2109.

9. Chen, Z., et al., Single-cell RNA sequencing highlights the role of inflammatory cancer-associated fibroblasts in bladder urothelial carcinoma. Nat Commun, 2020. 11(1): p. 5077.

10. Liu, Y., et al., Conserved spatial subtypes and cellular neighborhoods of cancer-associated fibroblasts revealed by single-cell spatial multi-omics. Cancer Cell, 2025. 43(5): p. 905–924 e6.

11. Strell, C., et al., Impact of Epithelial-Stromal Interactions on Peritumoral Fibroblasts in Ductal Carcinoma in Situ. Journal of the National Cancer Institute, 2019. 111(9).

12. Biffi, G., et al., Il1-induced Jak/STAT signaling is antagonized by TGFβ to shape CAF heterogeneity in pancreatic ductal adenocarcinoma. Cancer Discovery, 2019. 9(2): p. 282–301.

13. Mariathasan, S., et al., TGFβ attenuates tumour response to PD-L1 blockade by contributing to exclusion of T cells. Nature, 2018. 554(7693).

14. Tauriello, D.V.F., et al., TGFβ drives immune evasion in genetically reconstituted colon cancer metastasis. Nature, 2018. 554(7693): p. 538–543.

15. Giesen, C., et al., Highly multiplexed imaging of tumor tissues with subcellular resolution by mass cytometry. Nature Methods, 2014. 11(4): p. 417–422.

16. Tan, W.C.C., et al., Overview of multiplex immunohistochemistry/immunofluorescence techniques in the era of cancer immunotherapy. Cancer Commun (Lond), 2020. 40(4): p. 135–153.

17. Bankhead, P., et al., QuPath: Open source software for digital pathology image analysis. Scientific Reports, 2017. 7(1).

18. Ito, S., et al., Characterisation of colorectal cancer by hierarchical clustering analyses for five stroma-related markers. J Pathol Clin Res, 2024. 10(4): p. e12386.

19. Dominguez, C.X., et al., Single-cell RNA sequencing reveals stromal evolution into LRRC15+ myofibroblasts as a determinant of patient response to cancer immunotherapy. Cancer Discovery, 2020. 10(2): p. 232–253.

20. Elyada, E., et al., Cross-species single-cell analysis of pancreatic ductal adenocarcinoma reveals antigen-presenting cancer-associated fibroblasts. Cancer Discovery, 2019. 9: p. 1102–1123.

21. Xue, M., et al., Schwann cells regulate tumor cells and cancer-associated fibroblasts in the pancreatic ductal adenocarcinoma microenvironment. Nature Communications, 2023. 14(1).

22. Windhager, J., et al., An end-to-end workflow for multiplexed image processing and analysis. Nat Protoc, 2023. 18(11): p. 3565–3613.

23. Magness, A., et al., Deep cell phenotyping and spatial analysis of multiplexed imaging with TRACERx-PHLEX. Nature Communications, 2024. 15(1).

24. Bull, J.A., et al., MuSpAn: A Toolbox for Multiscale Spatial Analysis. 2024, Cold Spring Harbor Laboratory.

25. Jimenez-Sanchez, D., et al., NaroNet: Discovery of tumor microenvironment elements from highly multiplexed images. Med Image Anal, 2022. 78: p. 102384.

26. Wu, E., et al., ROSIE: AI generation of multiplex immunofluorescence staining from histopathology images. Nature Communications, 2025. 16(1).

