## Supplementary table and figures for "Development of a multiplex immunofluorescence panel to study heterogenous cancer-associated fibroblast subtypes with spatial resolution"

**Supplementary Table 1: A summary of the reagents used at comparable steps in the immunohistochemistry and immunofluorescence protocols.**

| Protocol step | IHC reagents list | IF reagents list |
|---|---|---|
| Tissue hydration | Xylene (Fisher Chemical, X/0250/17)<br>100%, 95%, 70% and 50% ethanol (VWR Chemical, 20821.330) | Xylene (Fisher Chemical, X/0250/17)<br>100%, 95%, 70% and 50% ethanol (VWR Chemical, 20821.330) |
| Antigen retrieval (AR) | IHC Antigen Retrieval Solution (Invitrogen low pH 00-4955-58 / high pH 00-4956-58) | 10X AR6 Buffer / 10X AR9 Buffer (Akoya Biosciences R600250ML / AR900250ML) |
| $H_20_2$ block | 3% hydrogen Peroxide ($H_2O_2$) Solution (Sigma Aldrich, 88597) | Novocastra Peroxidase Block (Leica Biosystems, RE7157) |
| Blocking | Protein Block Serum Free<br>(Dako, X0909) | Opal Antibody Diluent/Block (Akoya Biosciences ARD1001EA). |
| Primary antibody | Primary Antibody Diluent (Leica Biosystems, AR9352) | Opal Antibody Diluent/Block (Akoya Biosciences ARD1001EA). |
| Secondary antibody | SignalStain® Boost IHC Detection Reagent HRP (Cell Signaling Rabbit 8114S/ Mouse 8125S) | 1X Opal Anti-Ms +Rb HRP Kit (Akoya Biosciences ARH1001EA) |
| Detection | 3,3'-Diaminobenzidine (DAB) Chromogen (Novolink™ Max DAB polymer, Leica Biosystems, RE7270) | Opal 520 (Akoya Biosciences, OP-001001) diluted 1:150 in 1X Plus Automation Amplification Diluent (Akoya Biosciences, FP1609) |
| Nuclear counter stain | Haematoxylin<br>(Shandon Harris, REF6765004) | Spectral DAPI (Akoya Biosciences, FP1490) 1 drop diluted in 1mL TBST |
| Mount | Xylene based Cytoseal™<br>(Thermo Scientific, 8310-16) | Water based Immu-Mount<br>(Shandon, Immu-Mount™, 9990402) |

**Supplementary Table 2: Available positions for each CAF antigen in a multiplex panel.**

| Antigen | Available position |
|---|---|
| αSMA | 1,2,3,4,5 |
| FAP | 1 |
| PDGFRα | 3 |
| PDPN | 1,2,3,4,5 |

**Supplementary Table 3: A summary of the experimental design for each slide in the fluorescence minus one study.**

| Slide 1 | Slide 2 | Slide 3 | Slide 4 | Slide 5 | Slide 6 |
|---|---|---|---|---|---|
| FAP | *-2$^{nd}$ only-* | FAP | FAP | FAP | FAP |
| CD8 | CD8 | *-2$^{nd}$ only-* | CD8 | CD8 | CD8 |
| PDGFRα | PDGFRα | PDGFRα | *-2$^{nd}$ only-* | PDGFRα | PDGFRα |
| PDPN | PDPN | PDPN | PDPN | *-2$^{nd}$ only-* | PDPN |
| αSMA | αSMA | αSMA | αSMA | αSMA | *-2$^{nd}$ only-* |
| panCK | panCK | panCK | panCK | panCK | panCK |

**Supplementary Table 4: Threshold values for each single channel QuPath pixel classifier for evaluation of immunofluorescent staining.**

| Antigen | Opal | Threshold value |
|---|---|---|
| FAP | 690 | 0.4 |
| CD8 | 480 | 14 |
| PDGFRα | 570 | 0.5 |
| PDPN | 620 | 2.5 |
| αSMA | 520 | 0.9 |
| panCK | 780 | 0.5 |

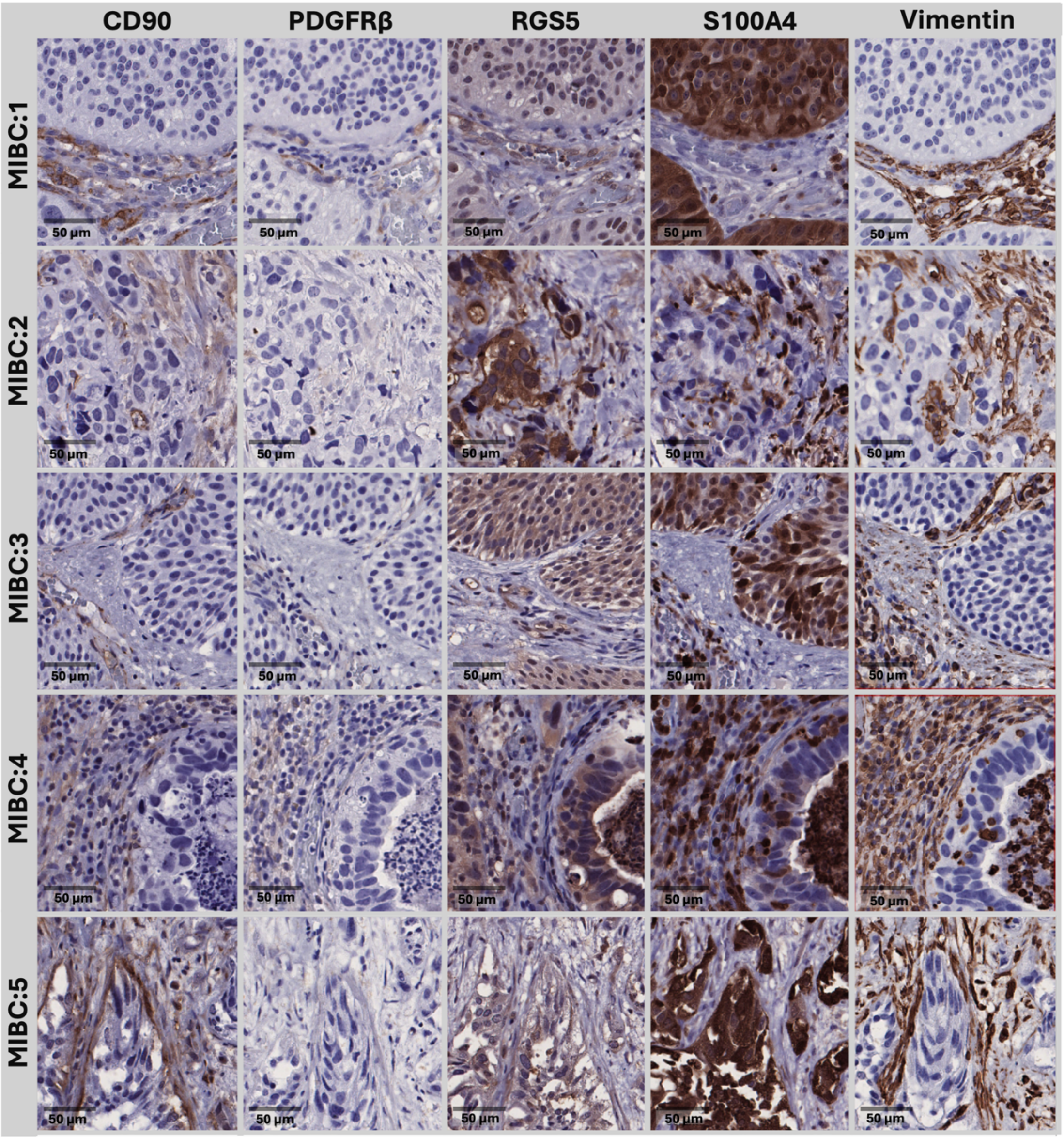


**Supplementary Figure 1: Immunohistochemistry conditions for antibodies considered but not included in downstream optimisation.**

Optimal IHC conditions shown in five MIBC samples that were stained with the CAF antigens that were considered but were not included in the final multiplex panel design. Each row represents a randomly selected region of tissue from a unique MIBC control, whilst each column highlights which antigen was stained for, positive staining is shown in brown (scale bar = 50 µM). CD90 ab92574 EPR3132 (Abcam), pH6.0, dilution 1:100. PDGFRβ ab69506 41G12 (Abcam), pH6.0, dilution 1:100. RGS5 ab196799 polyclonal (Abcam), pH6.0, dilution 1:100. S100A4 ab197896 EPR14639 (Abcam), pH9.0, dilution 1:2000. Vimentin ab92547 EPR3776 (Abcam), pH6.0, dilution 1:500.

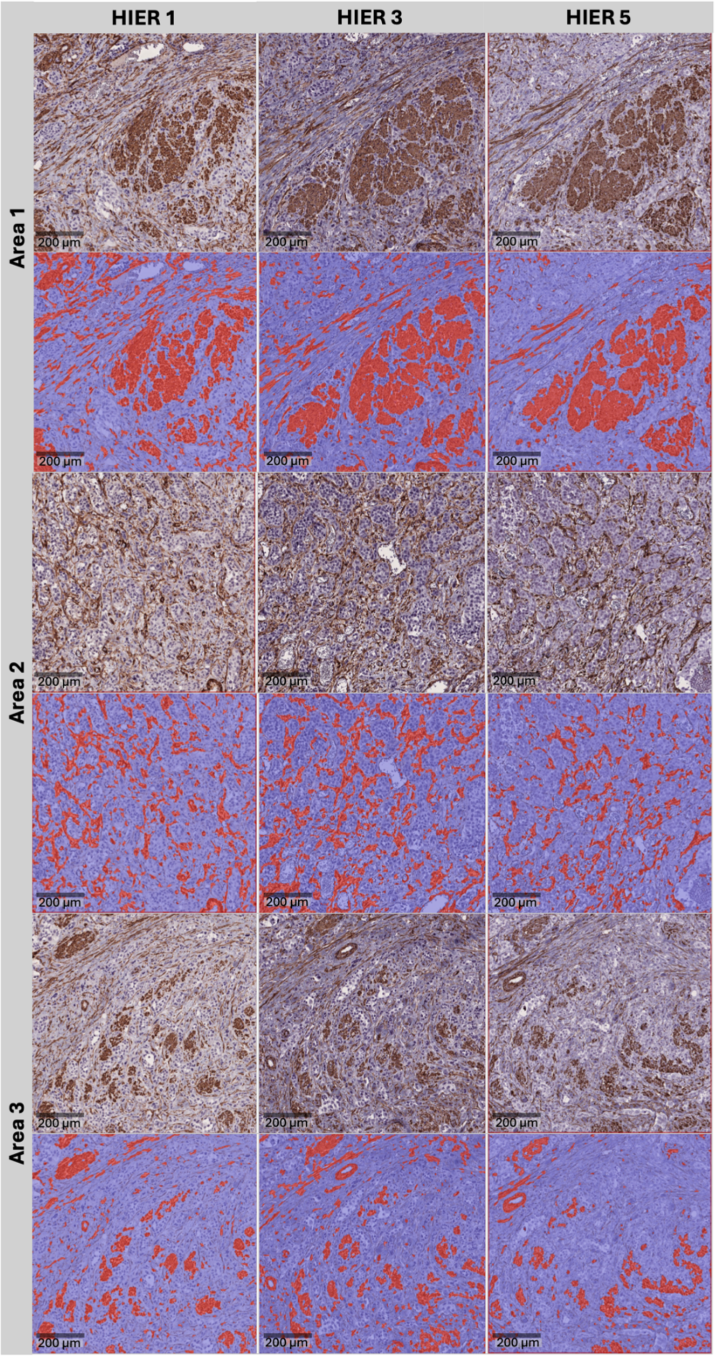


**Supplementary Figure 2: Assessment of αSMA antigen tolerance to heat-inactivation/ epitope retrieval.**

Example images show MIBC tissue stained with αSMA following 1, 3 and 5 rounds of HIER. Three distinct areas used for quantification in Figure 3B are displayed, the positive pixel detection of DAB staining is shown in red.

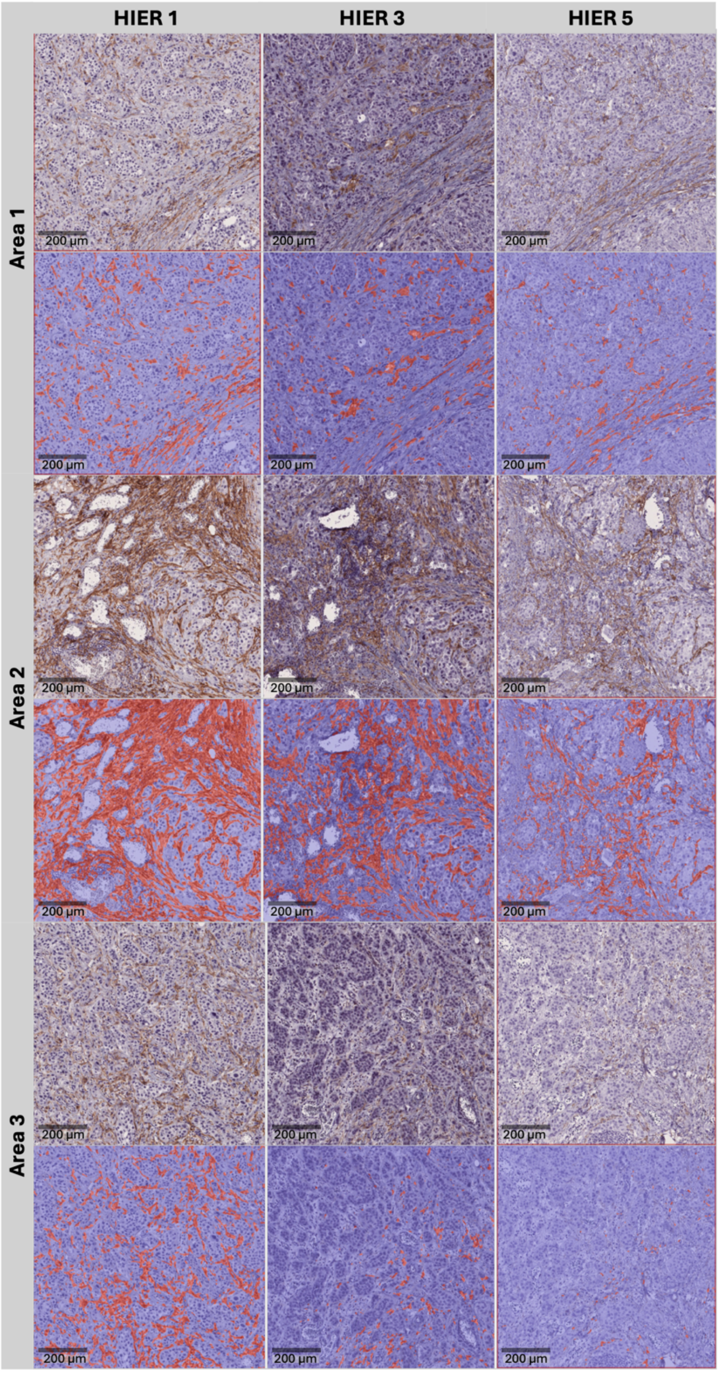


**Supplementary Figure 3: Assessment of FAP antigen tolerance to heat-inactivation/ epitope retrieval.**

Example images show MIBC tissue stained with FAP following 1, 3 and 5 rounds of HIER. Three distinct areas used for quantification in Figure 3B are displayed, the positive pixel detection of DAB staining is shown in red.

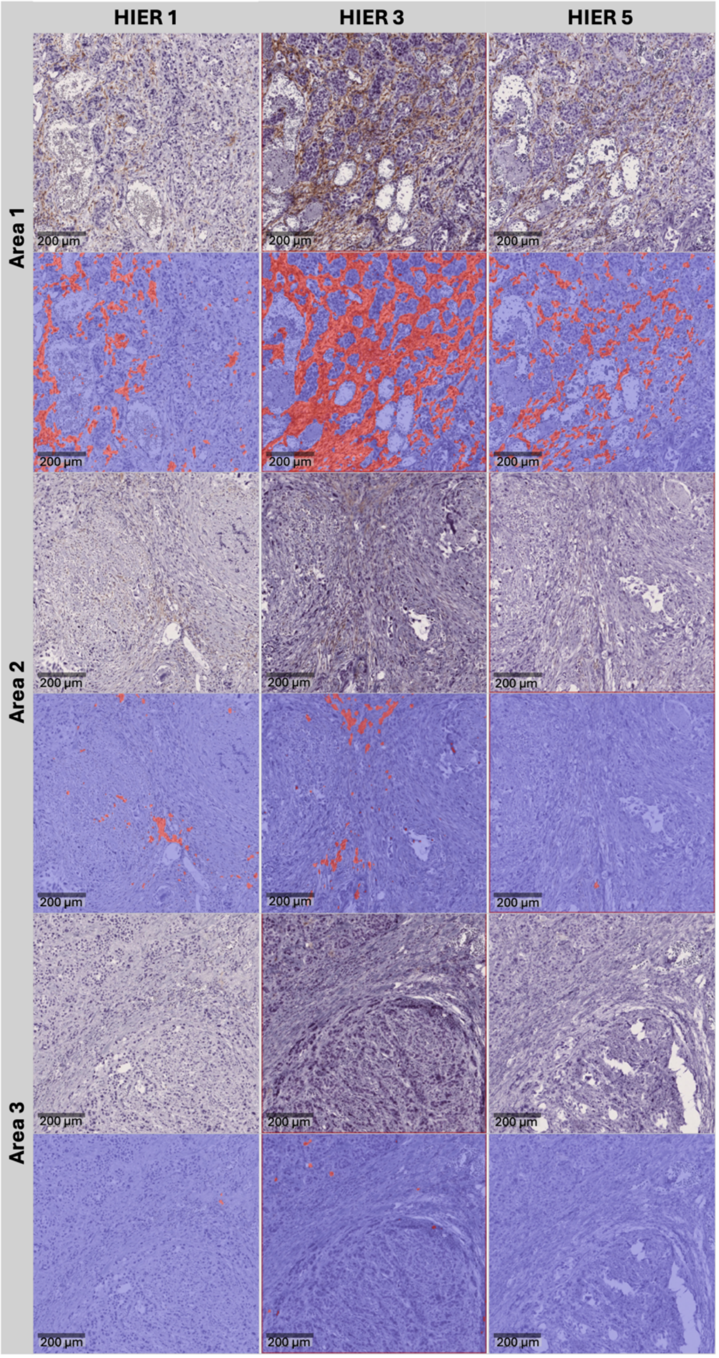


**Supplementary Figure 4: Assessment of PDGFRα antigen tolerance to heat-inactivation/ epitope retrieval.**

Example images show MIBC tissue stained with PDGFRα following 1, 3 and 5 rounds of HIER. Three distinct areas used for quantification in Figure 3B are displayed, the positive pixel detection of DAB staining is shown in red.

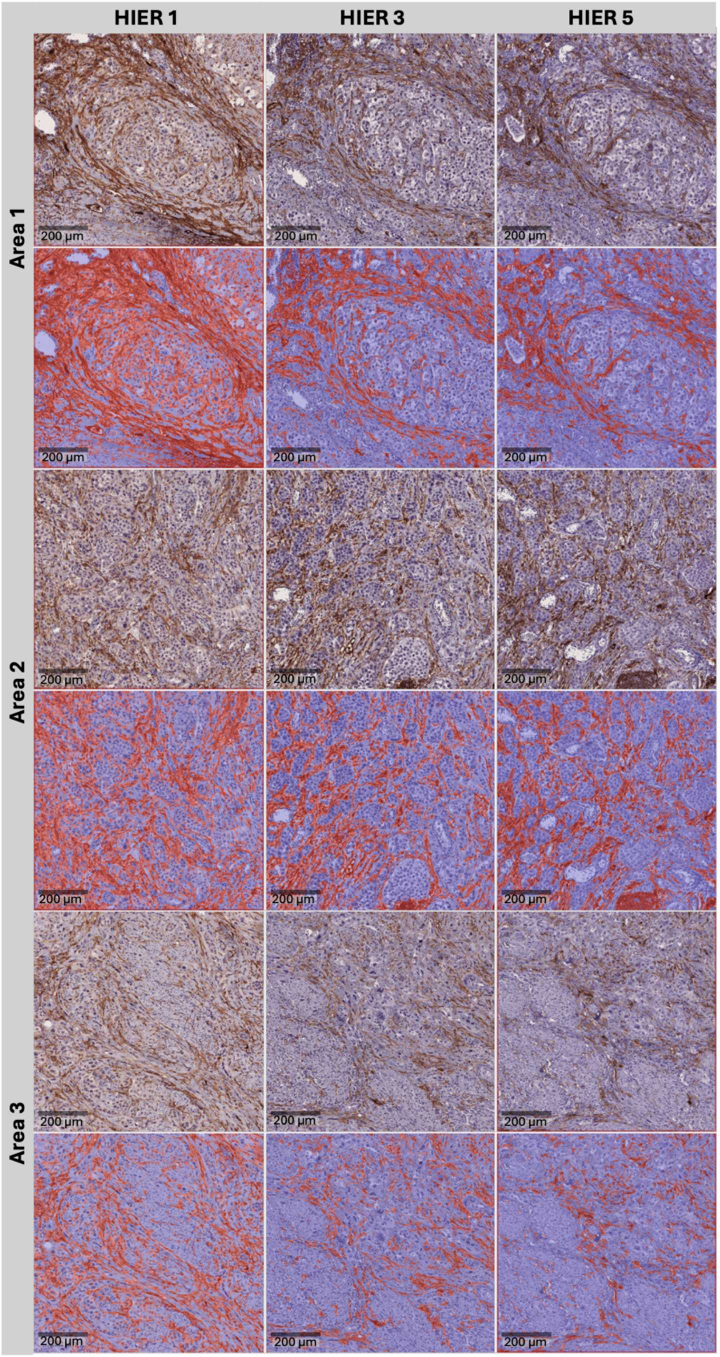


**Supplementary Figure 5: Assessment of PDPN antigen tolerance to heat-inactivation/ epitope retrieval.**

Example images show MIBC tissue stained with PDPN following 1, 3 and 5 rounds of HIER. Three distinct areas used for quantification in Figure 3B are displayed, the positive pixel detection of DAB staining is shown in red.

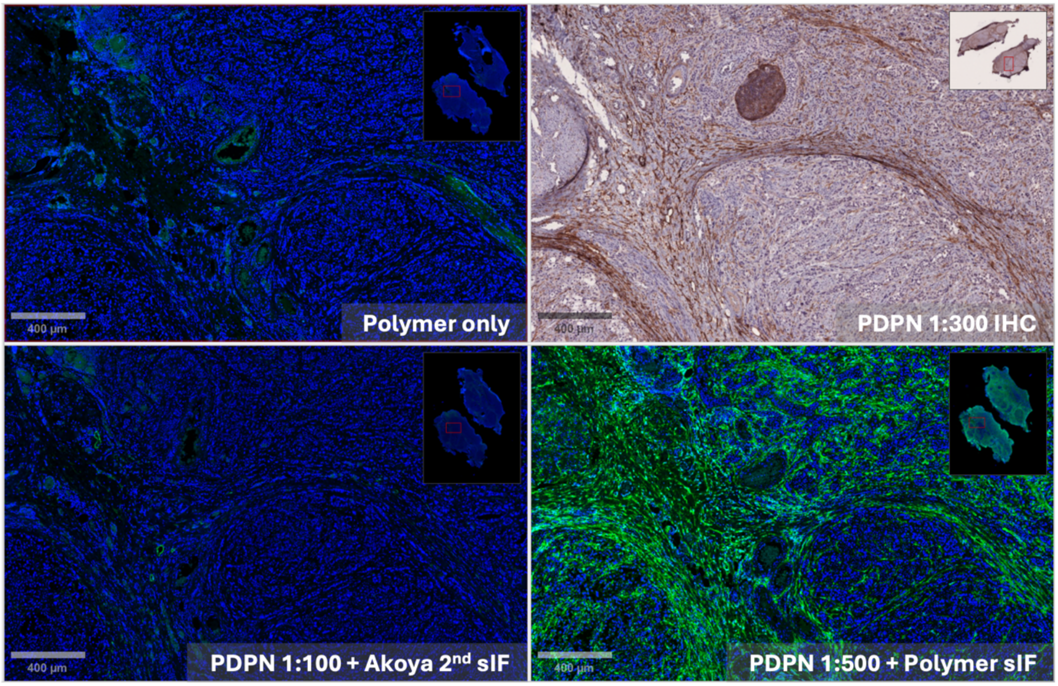


**Supplementary Figure 6: Qualitative assessment of PDPN single plex immunofluorescence staining with a polymer as an alternative secondary antibody.**

(Top left) An image of the polymer only immunofluorescence control, (top right) the PDPN IHC staining for reference, (lower left) PDPN sIF staining at 1:100 concentration with the Akoya secondary antibody for detection. (Lower right) PDPN sIF staining at 1:500 concentration with the Leica polymer for detection.

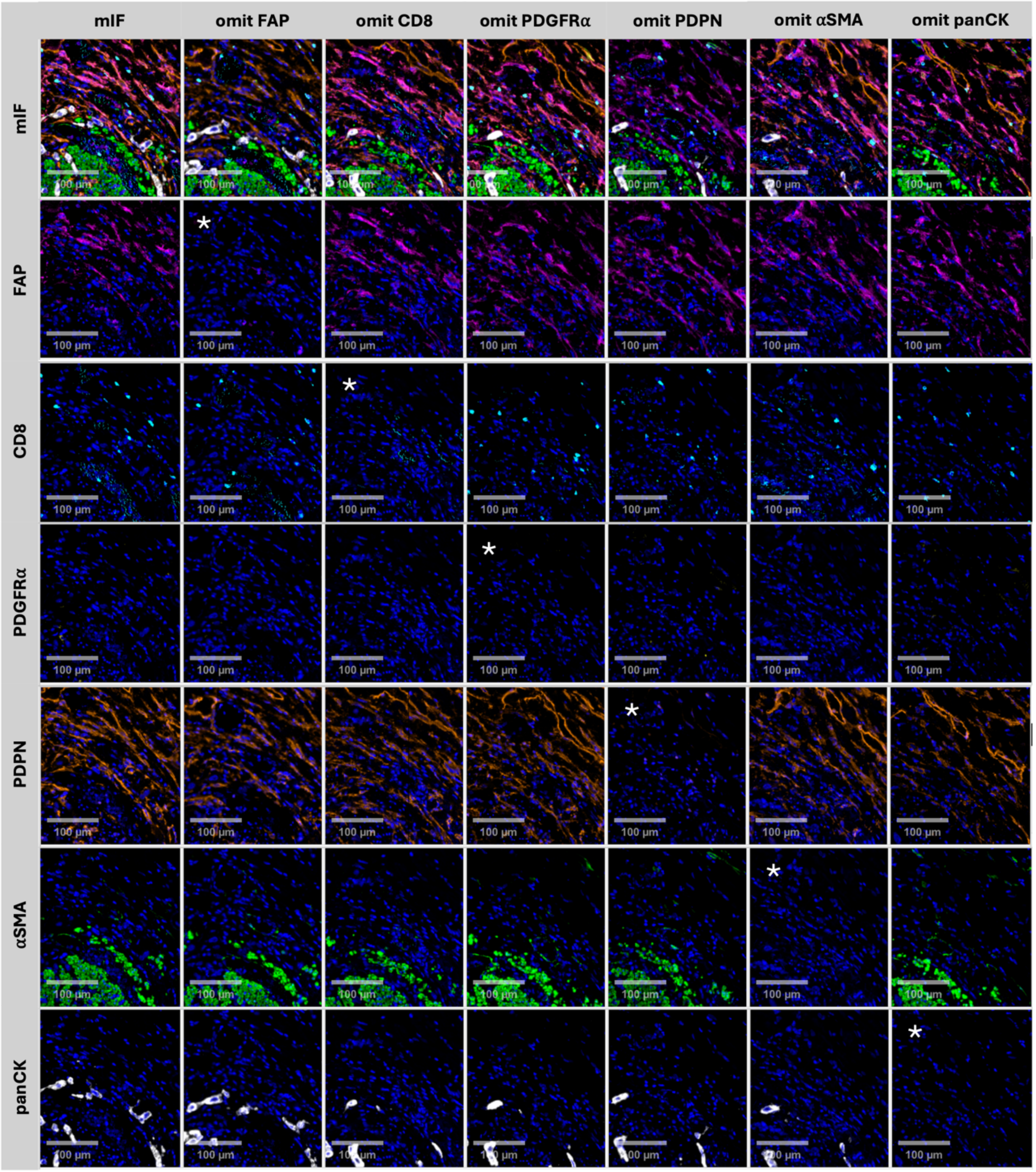


**Supplementary Figure 7: Confirmation of effective heat-inactivation in the multiplex panel using fluorescence minus one controls in a FAP-rich region.**

Example images of a FAP-rich region of MIBC tissue stained with the complete mIF panel or FMO controls where the primary antibody has been omitted, as indicated by an asterisk (scale bar = 100 µM). Each channel is shown individually to reveal the staining pattern of each antigen, the first column displays the full multiplex image for each slide.

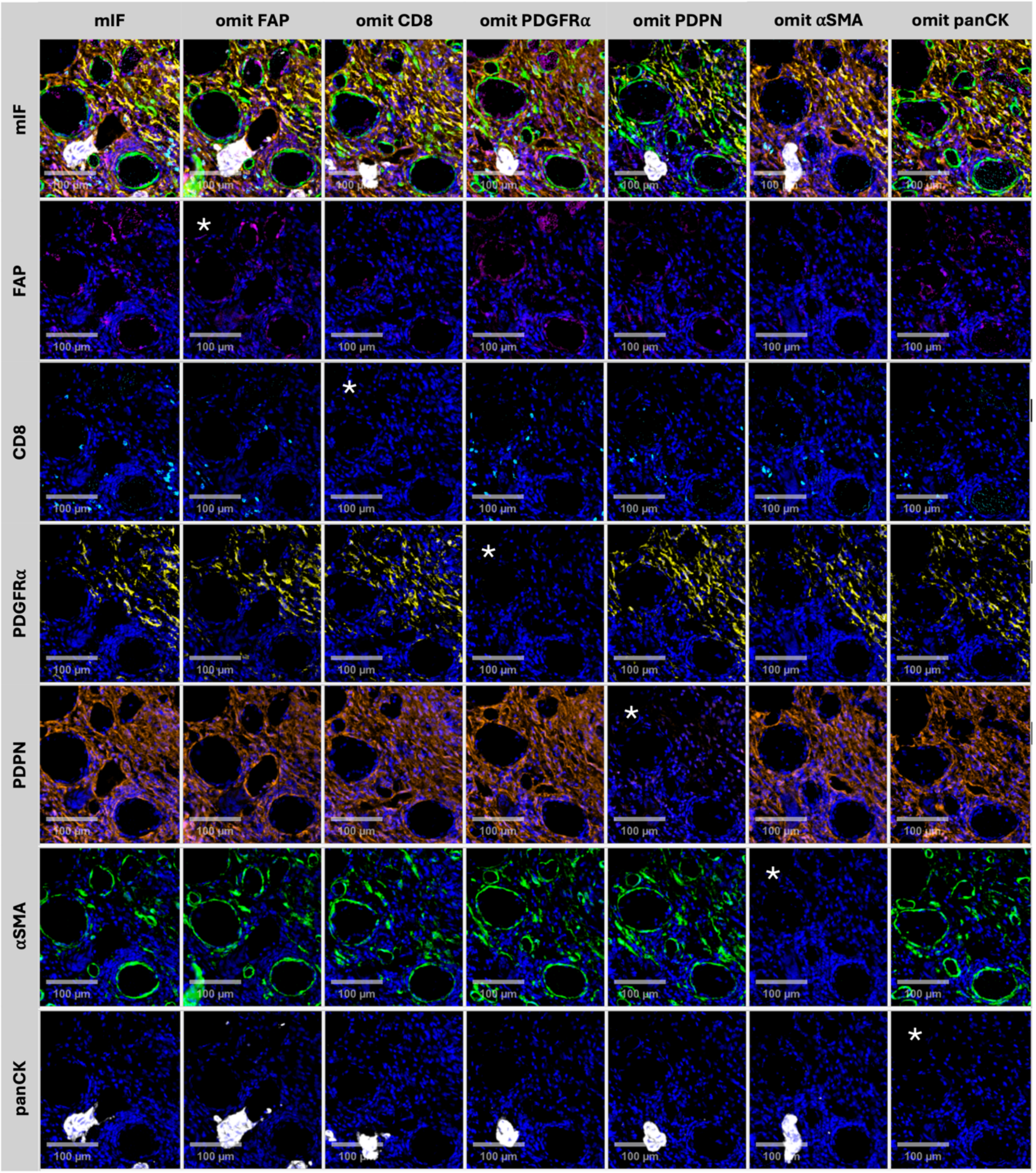


**Supplementary Figure 8: Confirmation of effective heat-inactivation in the multiplex panel using fluorescence minus one controls in a PDGFRα-rich region.**

Example images of a PDGFRα-rich region of MIBC tissue stained with the complete mIF panel or FMO controls where the primary antibody has been omitted, as indicated by an asterisk, (scale bar = 100 µM). Each channel is shown individually to reveal the staining pattern of each antigen, the first column displays the full multiplex image for each slide.

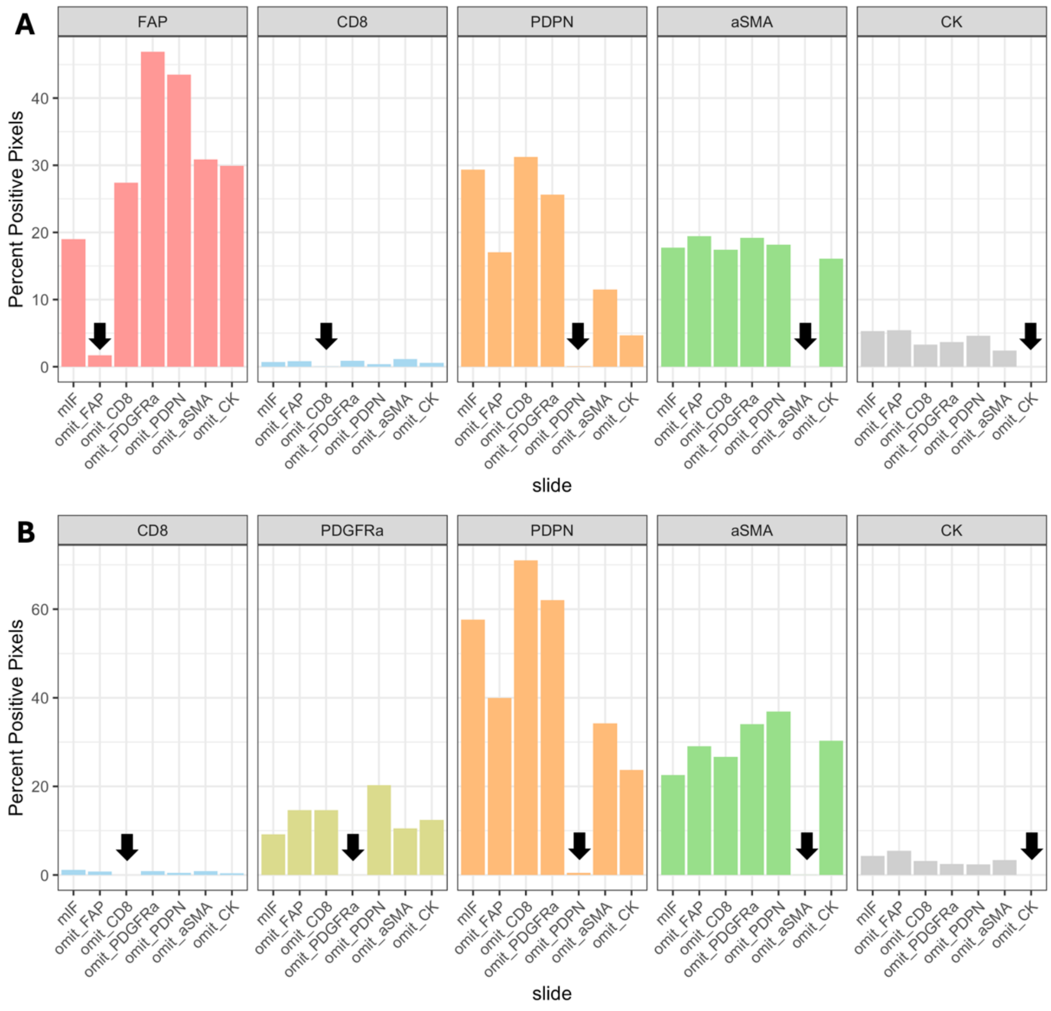


**Supplementary Figure 9: Quantification of fluorescence minus one experiment showing a depletion in signal that coincides with omission of each respective primary antibody.**

A single channel threshold was set for each marker in the multiplex panel, expression levels above the threshold contribute to the percentage of positive pixels in a given region of interest. Quantification of the FAP-rich region shown in Figure 7 shows effective depletion of FAP expression when the FAP primary antibody was omitted (A). To note, detection of auto fluorescent spots accounts for the small percentage detected in the 'omit FAP' slide. All other antibodies show effective clearance when the respective antibody is omitted (as indicated by the black arrowhead).

Quantification of the PDGFRα-rich region in Figure 8 shows effective clearance of PDGFRα expression when the PDGFRα primary antibody was omitted (B). Variation in the expression of the markers across slides is likely due to the variation in cell composition between serial sections and differences in the staining intensity.

**Supplementary files:**

json files for each IHC pixel classifier are provided in supplementary files.
